# Analysis of diffusion tensor imaging data from UK Biobank confirms dosage effect of 15q11.2 copy-number variation on white matter and shows association with cognition

**DOI:** 10.1101/2020.09.03.280859

**Authors:** Ana I. Silva, George Kirov, Kimberley M. Kendall, Mathew Bracher-Smith, Lawrence S. Wilkinson, Jeremy Hall, Magnus O. Ulfarsson, G. Bragi Walters, Hreinn Stefansson, Kari Stefansson, David E. J. Linden, Xavier Caseras

**Author notes:** Shared senior authorship (David E. J. Linden, Xavier Caseras). **Correspondent authors:** Ana I. Silva and Xavier Caseras, E-mail address, Address: Neuroscience and Mental Health Research Institute, MRC Centre for Neuropsychiatric Genetics and Genomics, Hadyn Ellis Building, Cathays, Cardiff, CF24 4HQ.

## Abstract

**Background:** Copy-number variations at the 15q11.2 BP1-BP2 locus are present in 0.5 to 1.0% of the population, and the deletion is associated with a range of neurodevelopmental disorders. Previously, we showed a reciprocal effect of 15q11.2 copy-number variation on fractional anisotropy, with widespread increases in deletion carriers. We aim to replicate and expand these findings, using a larger sample of participants (n=30,930), higher resolution imaging, and examining the implications for cognitive performance.

**Methods:** Diffusion tensor imaging measures from participants with no neurological/psychiatric diagnoses were obtained from the UK Biobank database. We compared 15q11.2 BP1-BP2 deletion (n=103) and duplication (n=119) carriers to a large cohort of control individuals with no neuropsychiatric copy-number variants (n=29,870). Additionally, we assessed how changes in white matter mediated the association between carrier status and cognitive performance.

**Results:** Deletion carriers showed increases in fractional anisotropy in the internal capsule and cingulum, and decreases in the posterior thalamic radiation, compared to both duplication carriers and controls (who had intermediate values). Deletion carriers had lower scores across cognitive tasks compared to controls, which were mildly influenced by white matter alterations. Reduced fractional anisotropy in the posterior thalamic radiation partially contributed to worse cognitive performance in deletion carriers.

**Conclusions:** This study, together with our previous findings, provides convergent evidence for a dosage-dependent effect of 15q11.2 BP1-BP2 on white matter microstructure. Additionally, changes in white matter were found to partially mediate cognitive ability in deletion carriers, providing a link between white matter changes in 15q11.2 BP1-BP2 carriers and cognitive function.

## Introduction

Several copy-number variants (CNVs) are associated with neurodevelopmental disorders (NDDs) in genome-wide studies, including intellectual disability (ID), autism spectrum disorder (ASD), epilepsy and schizophrenia (1–3). Altered white matter is common in many NDDs, and has been shown to mediate core cognitive deficits in schizophrenia (4,5). However, whether alterations in white matter microstructure are associated with these CNVs and can explain – at least partly – cognitive deficits in carriers of the risk variants, is not yet fully understood.

Among CNVs associated with NDDs, deletions and duplications at 15q11.2 are the most prevalent in humans, being present in 0.5% to 1.0% of the general population (6,7). 15q11.2 deletions are associated with developmental and motor delays (8), as well as increased susceptibility to attention deficit hyperactivity disorder (ADHD), ASD, schizophrenia, epilepsy (1,9) and congenital heart disease (10–12); whereas the pathogenicity of the corresponding duplication is less clear in population samples, where a significant risk for NDD has not been established (1,3) despite its link to neurodevelopmental phenotypes in clinical samples (13,14). Likewise, carriers of the 15q11.2 deletion unaffected by a diagnosed NDD, still show lower cognitive function than controls (10,15), as well as a higher prevalence of dyslexia and dyscalculia (6,16), whereas carriers of the 15q11.2 duplication perform similarly to controls on many cognitive tasks (15,16).

The 15q11.2 BP1-BP2 interval comprises four genes: nonimprinted in Prader-Willi/Angelman syndrome 1 (*NIPA1*), nonimprinted in Prader-Willi/Angelman syndrome 2 gene (*NIPA2*), cytoplasmic FMR1 interacting protein 1 (*CYFIP1*), and tubulin gamma complex associated protein 5 (*TUBGCP5*) (17). These genes are expressed in the central nervous system and have been individually associated with multiple disorders: *NIPA1* with autosomal-dominant hereditary spastic paraplegia (18), *NIPA2* with childhood absence epilepsy (19), *TUBHGCP5* with ADHD and obsessive-compulsive disorder (9), and *CYFIP1* with increasing susceptibility to ASD (20) and schizophrenia (21). A recent report investigated protein-protein interactions of the four genes in this region and found that their respective encoded proteins interact with each other, and that their predicted functions encompass crucial biological processes that are important for normal neuronal development, plasticity and function (22).

Using diffusion tensor imaging (DTI), we have previously demonstrated an association between 15q11.2 CNV dosage and altered white matter microstructure in an Icelandic sample (16). We found widespread increases of fractional anisotropy (FA) in deletion carriers relative to duplication carriers – with non-carrier controls showing intermediate values – and the largest effects were observed in the posterior limb of the internal capsule (23). The Icelandic gene pool is less heterogeneous than that of most European populations (24), facilitating the reduction of background noise caused by genetic variation (25), but arguably also raising concerns about the replicability of these findings in more genetically diverse, heterogeneous populations (26).

The aim of our study was to ascertain whether the above association between FA and 15q11.2 dosage was also present in a more heterogeneous European population like the UK, and whether FA mediates the known negative association between the 15q11.2 deletion and cognitive performance. To this end, we used a subsample of participants (~30 000) from the UK Biobank (www.ukbiobank.uk) for whom DTI-derived measures along with genetic data are available. Based on our previous findings, we hypothesized that bidirectional CNV dosage would lead to opposite changes in DTI measures in several white matter tracts, and that these changes could influence cognitive performance in deletion carriers.

## Methods and Materials

### Participants

A subsample of participants from the UK Biobank (www.ukbiobank.ac.uk) was used in this study. Ethical approval was granted by the North West Multi-Centre Ethics committee, and all subjects provided informed consent to participate in the UK Biobank project. Data were released to Cardiff University after application to the UK Biobank (project ref. 17044).

Only participants self-reporting white British or Irish descent and for whom genetic analysis confirmed European ancestry (27) were included (43,099 participants removed). Furthermore, in order to avoid confounding effects of disease, only participants with no personal history – based on self-reported diagnosis from a doctor at any assessment visit or existing hospital records – of neuropsychiatric disorders (i.e. schizophrenia, psychosis, ASD, dementia or ID) or medical/neurological conditions that could impact white matter (i.e. alcohol or other substance dependency, Parkinson’s, Alzheimer’s, multiple sclerosis or other neurodegenerative conditions) were selected (43,290 participants removed). After applying these exclusions, 416,253 participants remain.

### Genotyping, CNV calling and CNV quality control

DNA extraction and processing workflow are described at https://biobank.ctsu.ox.ac.uk/crystal/crystal/docs/genotyping_sample_workflow.pdf. CNV calling was performed by Kendall *et al.* (28), and quality control parameters are briefly explained in Supplemental Methods. Carriers of CNVs at the 15q11.2 BP1-BP2 locus, and participants with no neurodevelopmental CNVs (NoCNV) were selected. For the NoCNV group, we selected participants that carried none of the 93 CNVs that have previously been associated with NDDs (3,29,30). The 15q11.2 BP1-BP2 interval was manually inspected to confirm that included the key genes within the region (Table S1). None of the 15q11.2 BP1-BP2 CNV carriers included in this study carried any other large CNVs. Altogether, we found 1519 15q11.2 BP1-BP2 deletion carriers, 1833 duplication carriers, and 370,289 NoCNV carriers in the remaining sample after exclusions and quality control.

### Diffusion tensor imaging data

We used standard DTI measures made available by the UK Biobank. Detailed brain imaging protocols can be found in the Biobank brain MRI documentation (http://biobank.ctsu.ox.ac.uk/crystal/crystal/docs/brain_mri.pdf). DTI data were acquired using a multishell approach with two b-values (b=1000 and 2000 s/mm^2^). For each diffusion-weighted shell, 50 diffusion-encoding directions were acquired. Tensor fitting utilizes the b=1000 s/mm^2^ data, leading to the generation of FA, axial diffusivity (AD), radial diffusivity (RD), and mean diffusivity (MD) maps. The DTI maps were used in Tract-Based Spatial statistics (TBSS) processing, and TBSS-derived measures were computed by averaging the skeletonized images of each DTI map within a set of 48 standard-space tract masks defined by the JHU white matter atlas (ICBM-DTI-81) (31).

DTI data were available for 30,930 participants in the UK Biobank; of those 103 were 15q11.2 BP1-BP2 deletion carriers, 119 were 15q11.2 BP1-BP2 duplication carriers, and 29,870 were NoCNV carriers. Participants were aged between 40 and 70 years old, and the numbers of females and males were similar in each group. Demographic information is described in Table 1.

**Table 1.**
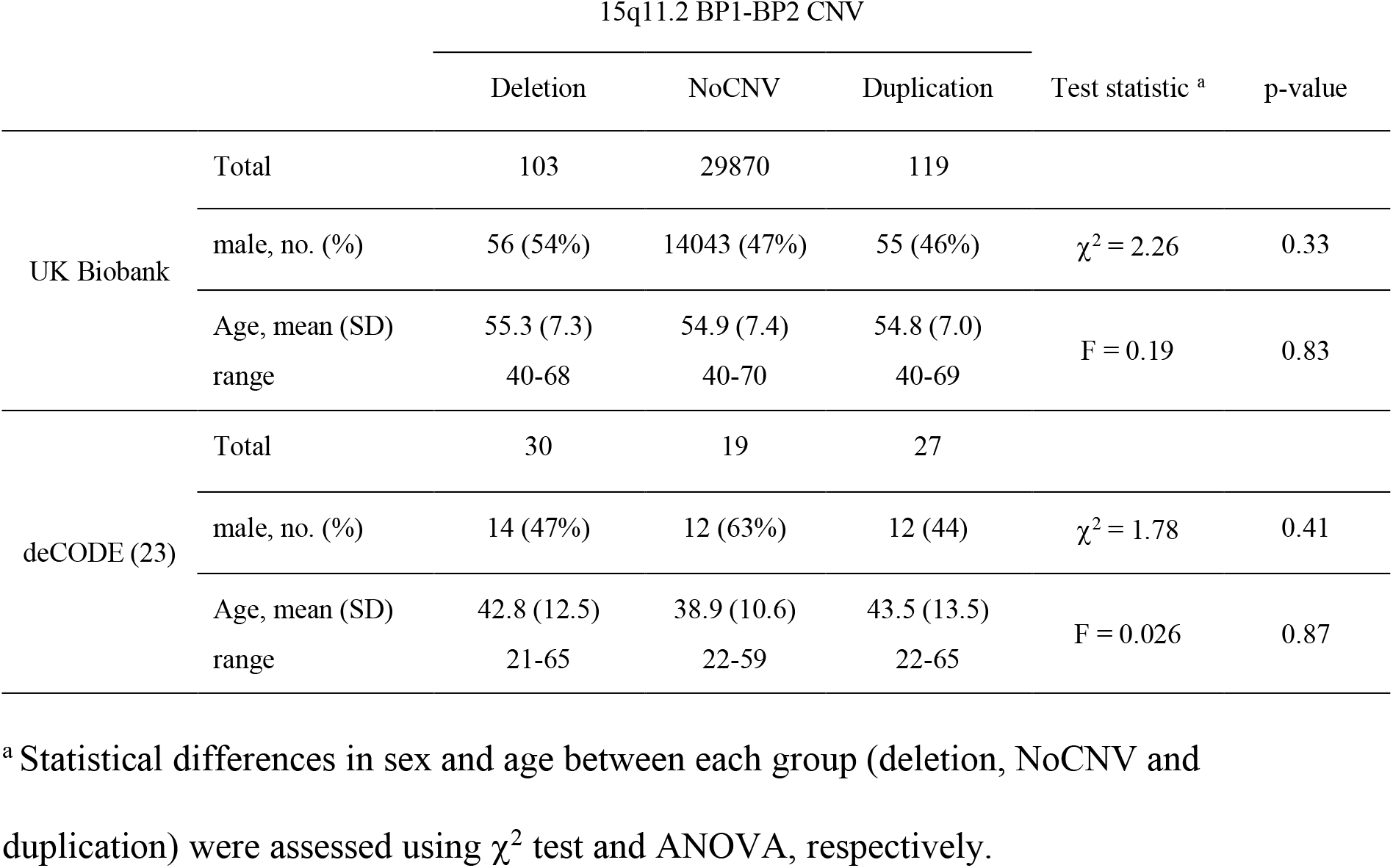
Demographic characteristics of individuals with neuroimaging data available from the UK Biobank, after exclusion of individuals with a known brain disorder, and after quality control of genotyping data (see methods for exclusion criteria). Details from the Icelandic sample, used in our previous study, are shown for comparison. **Abbreviations used:** CNV, copy-number variants; NoCNV, no pathogenic CNVs.

|  |  | 15q11.2 BP1-BP2 CNV |  |  | Test statistic <sup>a</sup> | p-value |
| --- | --- | --- | --- | --- | --- | --- |
|  |  | Deletion | NoCNV | Duplication |  |  |
| UK Biobank | Total | 103 | 29870 | 119 |  |  |
| | male, no. (%) | 56 (54%) | 14043 (47%) | 55 (46%) | $\chi^2 = 2.26$ | 0.33 |
|  | Age, mean (SD) | 55.3 (7.3) | 54.9 (7.4) | 54.8 (7.0) | F = 0.19 | 0.83 |
|  | range | 40-68 | 40-70 | 40-69 |  |  |
| deCODE (23) | Total | 30 | 19 | 27 |  |  |
| | male, no. (%) | 14 (47%) | 12 (63%) | 12 (44) | $\chi^2 = 1.78$ | 0.41 |
|  | Age, mean (SD) | 42.8 (12.5) | 38.9 (10.6) | 43.5 (13.5) | F = 0.026 | 0.87 |
|  | range | 21-65 | 22-59 | 22-65 |  |  |

<sup>a</sup> Statistical differences in sex and age between each group (deletion, NoCNV and duplication) were assessed using $\chi^2$ test and ANOVA, respectively.

To avoid the potential effect of extreme values that could have resulted from data quality or processing problems, outlier values of FA, AD, RD and MD – defined as values ± 2.5 standard deviations from each group mean – in each white matter tract were removed from the analyses (for that particular white matter tract).

### Cognitive data

Participants of UK Biobank also underwent a series of cognitive tests. We evaluated the performance on seven cognitive tasks – the pairs matching, reaction time, fluid intelligence, digit span, symbol digit substitution, and trail making A and B tasks. Data transformations of cognitive measures taken were performed after assessment of normality, following the approach of Kendall *et al.* (28). All cognitive measures were transformed so that lower values represented poorer performance. Table 3 describes the sample sizes used for each task in our neuroimaging sample. Since cognitive data are available for many more participants in the UK Biobank than those with neuroimaging data, we also report an extended analysis considering the full sample in Table S3. Details about each cognitive task can be found in Supplemental Methods.

### Statistical analyses

The mean TBSS-derived measures (FA, AD, RD and MD) from 30 white matter tracts were considered for analyses (Table 2). We focused our analyses on tracts that have previously been associated with psychiatric disorders (32). Statistical analyses were performed in R *version 3.6.3* (R Foundation).

**Table 2.** The 30 white matter tracts selected for this study and the abbreviations.

| White matter tracts | Abbreviations |
| --- | --- |
| Genu Corpus Callosum | GCC |
| Body Corpus Callosum | BCC |
| Splenium Corpus Callosum | SCC |
| Fornix | Fornix |
| Corticospinal tract - Right and left | CST_R; CST_L |
| Anterior Limb of the Internal Capsule - right and left | ALIC_R; ALIC_L |
| Posterior Limb of the Internal Capsule - right and left | PLIC_R; PLIC_L |
| Anterior Corona Radiata - right and left | ACR_R; ACR_L |
| Superior Corona Radiata - right and left | SCR_R; SCR_L |
| Posterior Corona Radiata - right and left | PCR_R; PCR_L |
| Posterior Thalamic Radiation - right and left | PTR_R; PTR_L |
| Sagittal Stratum (includes inferior longitudinal fasciculus) - right and left | SS_R; SS_L |
| External Capsule - right and left | EC_R; EC_L |
| Cingulum (cingulate gyrus portion) - right and left | C_CG_R; C_CG_L |
| Cingulum (hippocampus portion) - right and left | C_HIP_R; C_HIP_L |
| Superior Longitudinal Fasciculus - right and left | SLF_R; SLF_L |
| Uncinate Fasciculus - right and left | UF_R; UF_L |

To evaluate group differences in TBSS-derived and cognitive measures, we performed an ANOVA, including age, sex and handedness as covariates. For brain measures we also included brain size (total grey matter + white matter volumes) as a covariate. Following this, *post hoc* pairwise comparisons were performed to measure differences between groups (*deletion vs NoCNV, duplication vs NoCNV,* and *deletion vs duplication).* We used the Benjamini and Hochberg (BH) false discovery rate (FDR, q=0.05) to account for multiple testing (33), considering the relationship/high degree of concordance between the various white matter tracts and within the DTI metrics, as well as differing cognitive measures. The interaction between copy-number and age, and between copy-number and sex was also assessed and results are shown in Supplemental Findings and in Tables S4-S7.

Cohen’s d effect sizes were calculated, comparing *deletion vs NoCNV* and *duplication vs NoCNV*, and plotted using diverging bars. Adjusted values for each group were used, regressing out the effects of age, sex, handedness and brain volume using linear regression. Cohen classified effect sizes as negligible (d<0.2), small (0.2<d<0.5), medium (0.5<d<0.8), and large (d>0.8) (34).

To assess the degree of overlap between our current findings, in the UK Biobank sample, and our previous findings, in the Icelandic sample (23), we plotted effect sizes from both samples using forest plots. We recalculated the effect sizes from the Icelandic sample using adjusted values for age, sex and brain volume, and only white matter tracts showing group differences in either study are shown (Figures S1 and S2).

Mediation analysis was performed to test the hypothesis that 15q11.2 BP1-BP2 CNV effects on cognition are mediated by white matter abnormalities. Tracts showing a significant association between FA and carrier status were considered, as well as cognitive tasks that were significantly affected in carriers, when compared to NoCNV carriers. Linear regression was used to look at overall effects of white matter on cognitive tasks (including all deletion- and duplication carriers). Mediation analysis was conducted using the mediation package *version 4.4.7* in R, which uses structural equation modelling. We report the proportion of the total effect of copy-number on cognitive performance mediated by FA, with p-values calculated through quasi-Bayesian approximation using 5000 simulations. Age, sex, handedness and brain volume were included as covariates. FDR correction was again applied to account for multiple testing.

## Results

### Group differences on TBSS-derived measures

Most differences were seen in the contrast between deletion- and duplication carriers, confirming the reciprocal effect of 15q11.2 BP1-BP2 dosage previously reported on white matter microstructure (23).

15q11.2 BP1-BP2 deletion carriers showed increased FA relative to duplication carriers in ALIC_L, PLIC_R, PLIC_L, C_HIP_R and C_HIP_L, and decreased FA in fornix and PTR_R (Figure 1 and Table S2 for descriptive statistics). Deletion carriers also showed increased FA in ALIC_L, PLIC_L and C_HIP_L, and decreased FA in PTR_R, when compared to NoCNV carriers. Additionally, deletion carriers showed significant decreases in MD in BCC and UF_L, and significant decreases in AD in BCC and SCC, when compared to NoCNV carriers. 15q11.2 BP1-BP2 duplication carriers only showed reduced FA in the C_HIP_R compared to NoCNV carriers.

**Figure 1.**
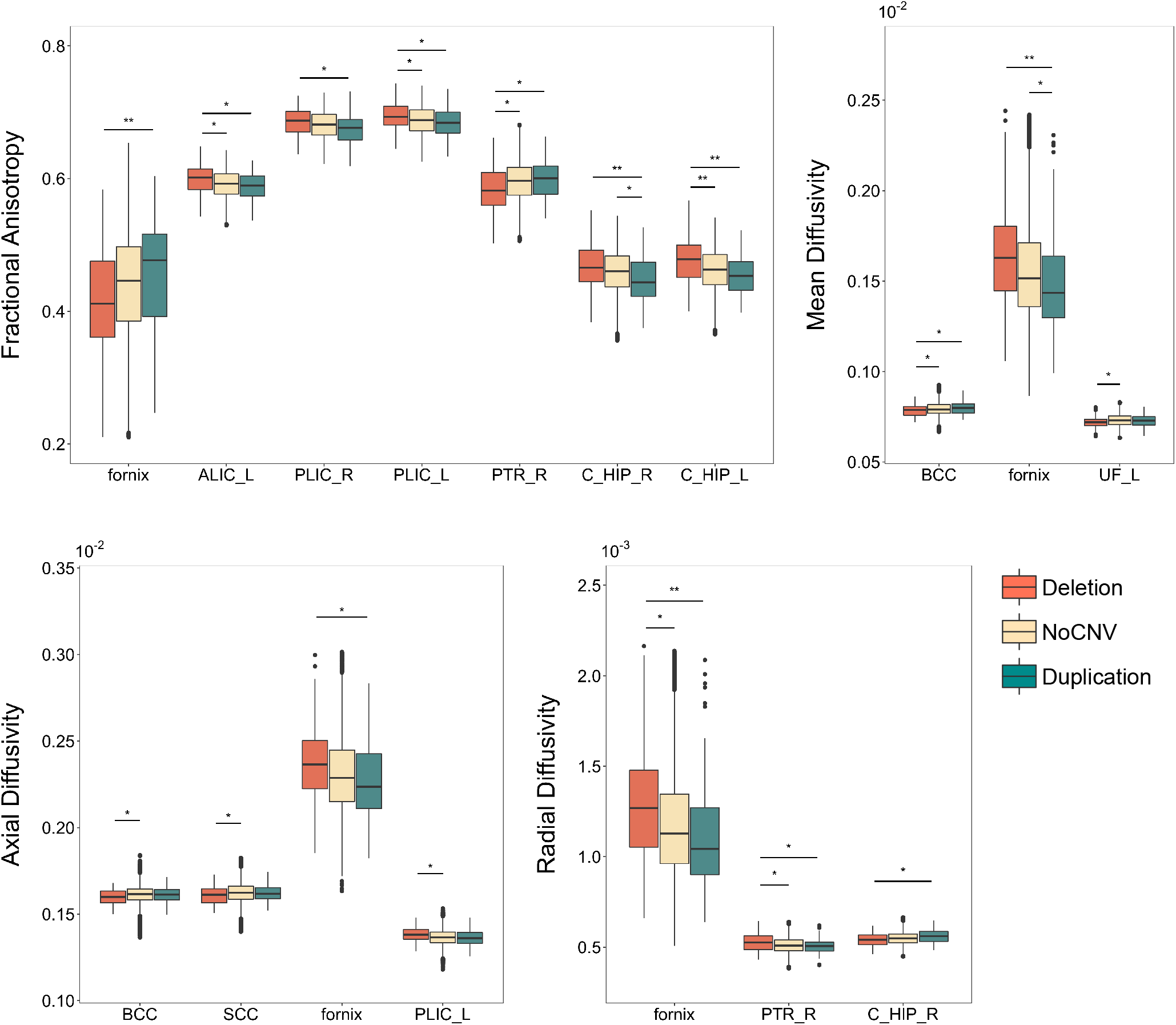
Boxplots showing the effects of 15q11.2 BP1-BP2 copy-number variation on TBSS-derived measures. Group differences between deletion (n=103), NoCNVs (n=29,870) and duplication (n=119) carriers were assessed with a linear regression followed by *post hoc* pairwise comparisons. Here, only white matter tracts showing significant group differences after FDR correction for multiple comparisons, are shown. Statistics are documented in Table S2.

The effect sizes for FA were overall small (Cohen’s d<0.5) when comparing CNV carriers to NoCNV carriers but were higher when comparing deletion- to duplication carriers, where medium effect sizes (Cohen’s d>0.5) were found in C_HIP_R, and C_HIP_L, underlining the reciprocal CNV dosage effect in these regions (Table S2). For AD, RD and MD, a medium effect size was also seen in the fornix, when comparing deletion- to duplication carriers. Diverging bar plots showing effect sizes for all thirty white matter tracts considered, are shown in Figures S3-S6.

### Cognitive performance in 15q11.2 CNV carriers

Deletion- and duplication carriers were affected differently regarding cognitive performance, when compared to NoCNV carriers, in our neuroimaging sample: deletion carriers showed poorer performance on reaction time, fluid intelligence, symbol substitution and trail making B tasks, whereas duplication carriers achieved a similar level of performance as NoCNV carriers (Table 3). This was also true when considering all participants with cognitive data (Table S3), where deletion carriers showed poorer performance in pairs matching, reaction time, fluid intelligence, digit span, symbol substitution and trail making B tasks; duplication carriers showed poorer performance only in pairs matching task, and no effects on other tasks.

**Table 3.** Effects of 15q11.2 BP1-BP2 copy-number variation on seven cognitive tasks from the UK Biobank. Only individuals that were used in the neuroimaging analysis were considered. Group differences were assessed using an ANOVA followed by *post hoc* pairwise comparisons, and corrected using FDR correction. Both uncorrected and corrected (BH corr.) p-values are shown. **Abbreviations used:** Del, deletion; Dup, duplication; NoCNV, no pathogenic copy-number variants.

|  | 15q11.2 BP1-BP2 CNV |  |  | Deletion vs NoCNV |  |  | Duplication vs NoCNV |  |  | Dosage |  |
| --- | --- | --- | --- | --- | --- | --- | --- | --- | --- | --- | --- |
|  | Del, No. | NoCNV, No. | Dup, No. | Cohen's d (SE) | p-value | p-value (BH corr.) | Cohen's d (SE) | p-value | p-value (BH corr.) | F statistic | p-value |
| Pairs matching | 103 | 29497 | 118 | -0.06 (0.19) | 0.54 | 0.63 | 0.002 (0.18) | 0.98 | 0.98 | $F_{(2,29710)} = 0.23$ | 0.79 |
| Reaction time | 103 | 29815 | 119 | -0.34 (0.19) | 0.0006 | **0.007** | 0.16 (0.18) | 0.09 | 0.17 | $F_{(2,30029)} = 7.40$ | 0.0006 |
| Fluid intelligence | 90 | 27532 | 114 | -0.33 (0.21) | 0.0018 | **0.01** | -0.06 (0.18) | 0.49 | 0.6 | $F_{(2,27728)} = 4.86$ | 0.008 |
| Digit span | 72 | 20433 | 87 | -0.28 (0.23) | 0.02 | 0.05 | -0.11 (0.21) | 0.32 | 0.48 | $F_{(2,20584)} = 3.07$ | 0.05 |
| Symbol substitution | 62 | 15600 | 59 | -0.36 (0.25) | 0.005 | **0.02** | 0.10 (0.26) | 0.44 | 0.57 | $F_{(2,15713)} = 4.50$ | 0.01 |
| Trail making A | 52 | 13966 | 52 | -0.29 (0.27) | 0.04 | 0.09 | -0.07 (0.27) | 0.59 | 0.65 | $F_{(2,14062)} = 2.69$ | 0.07 |
| Trail making B | 52 | 13966 | 52 | -0.40 (0.27) | 0.004 | **0.02** | 0.11 (0.27) | 0.42 | 0.57 | $F_{(2,14062)} = 5.15$ | 0.006 |

As described above, four white matter tracts (ALIC_L, PLIC_L, PTR_R and C_HIP_L) showed FA changes in deletion carriers, and one tract (C_HIP_R) in duplication carriers, when compared to NoCNV carriers. Therefore, the effects of these tracts on fluid intelligence, reaction time, symbol substitution and trail making B task performance were tested through regression analysis. FA variation in all these tracts were overall significantly associated with cognitive task performance, where increases in FA were associated with better performance, with the exception of PLIC_L where no significant associations were found for any of the tasks (Table 4).

**Table 4.**
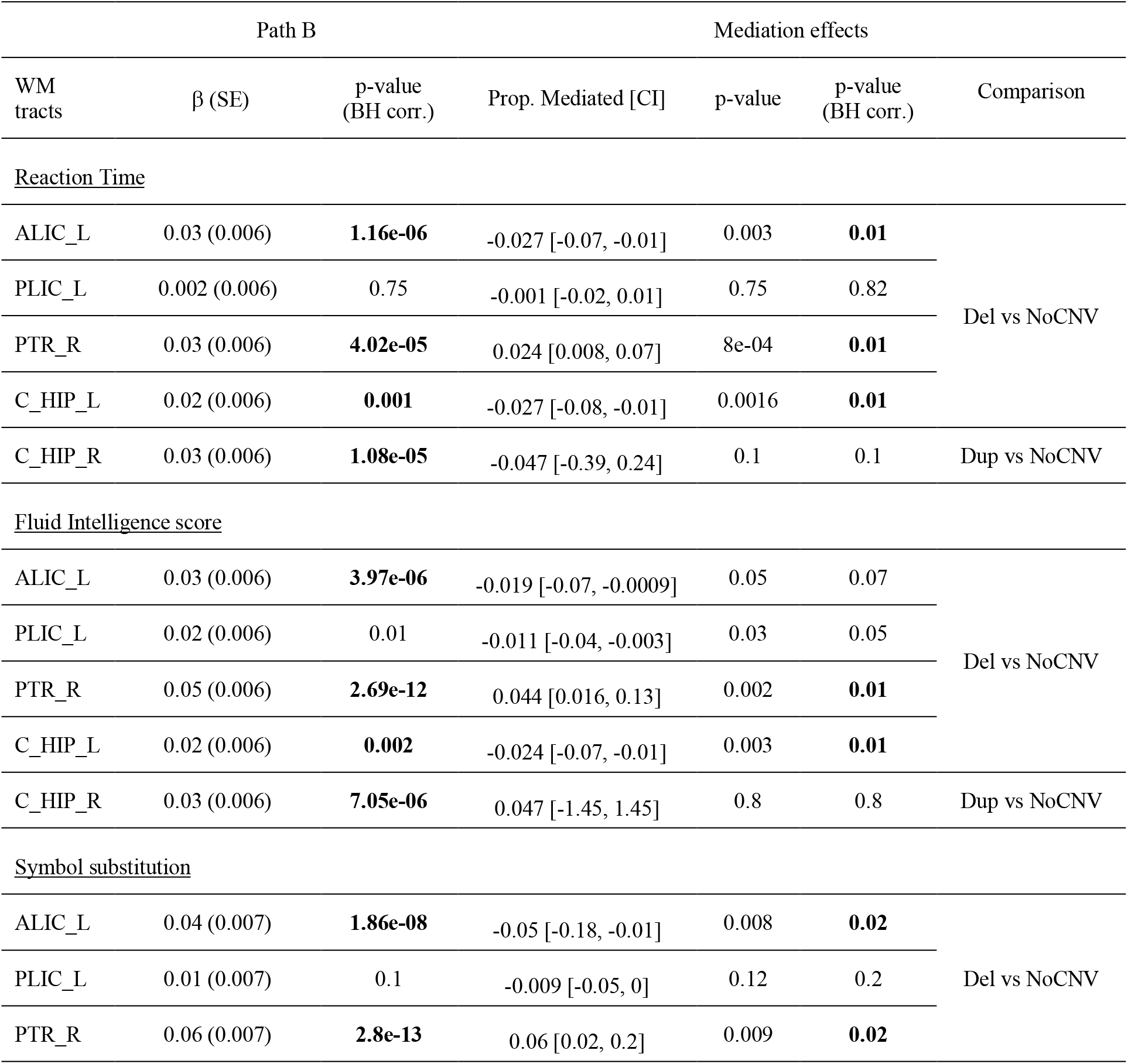
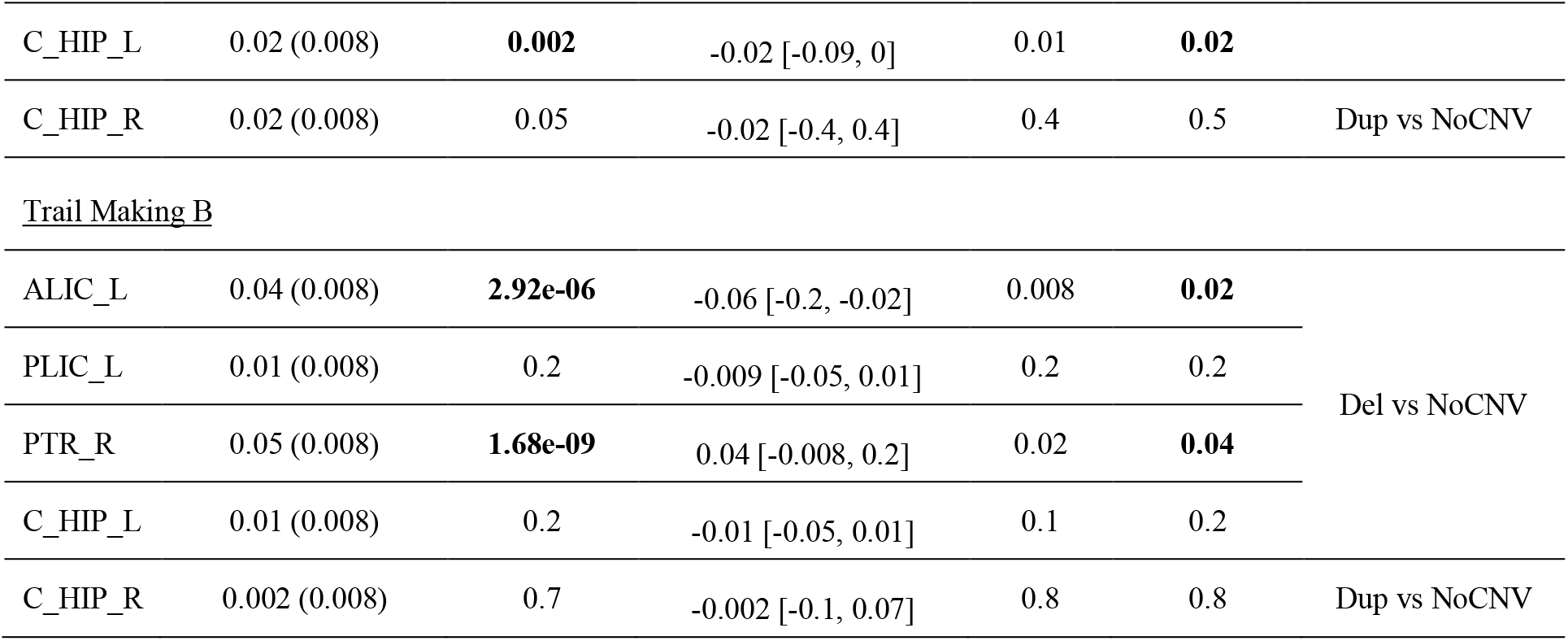
Mediation analysis showing the proportion of the total effect of 15q11.2 BP1-BP2 CNV on reaction time, fluid intelligence, symbol substitution and trail making B, mediated by fractional anisotropy (FA) from different white matter tracts. Negative proportions indicate opposite signs between the mediator (FA) and the total effect (CNV effect on cognitive measure). The linear regression (Path B) shows the overall effects of white matter on cognition (including all deletion- and duplication carriers). Both uncorrected and corrected (BH corr.) p-values are shown. **Abbreviations used:** WM, white matter; Del, deletion; Dup, duplication; NoCNV, no pathogenic copy-number variants.

We found that ALIC_L, PTR_R and C_HIP_L partially mediated cognitive performance in deletion carriers. Decreased FA in PTR_R was found to partially mediate cognitive performance on all four cognitive tasks, accounting for between 2.4% and 6% of the poorer performance of deletion carriers (Table 4). Increased FA in ALIC_L in deletion carriers was associated with higher scores in reaction time, symbol substitution and trail making B tasks, removing 2.7%, 5%, and 6% of the total effect of the CNV on each task respectively. Similarly, increased FA in C_HIP_L removed 2.7%, 2.4% and 2% of the total effect of the CNV on reaction time, fluid intelligence score and symbol substitution, respectively.

## Discussion

This is the largest study to date investigating the effects of the 15q11.2 BP1-BP2 CNV on white matter microstructure, as well as the first study examining how these effects are associated with cognitive ability. Using a large sample from UK Biobank, we found significant CNV dosage effects on TBSS-derived measures, with NoCNV carriers placed between deletion- and duplication carriers in most tracts, suggesting a dosage-dependent effect on white matter. We found more prominent differences between deletion- and NoCNV carriers than between duplication- and NoCNV carriers. These results replicate our previous findings of reciprocal white matter changes in 15q11.2 BP1-BP2 CNV carriers in an Icelandic sample, with larger effect sizes in deletion-than duplication carriers (23). Furthermore, we showed that deletion carriers have poorer cognitive performance, which is partially mediated by FA measures.

Previously, in the Icelandic sample, we found increased FA in deletion carriers in the left inferior longitudinal fasciculus (ILF_L), PCR_L, PTR_R, C_CG_L, ALIC_L, PLIC_R and PLIC_L, compared to duplication carriers, but no significant differences were found between carriers and non-carriers (23). The UK Biobank data replicated the copy-number effects on ALIC_L, PLIC_R and PLIC_L, where significant increases in FA were seen in deletion carriers, compared to duplication carriers. Additionally, significant differences between deletion- and NoCNV carriers were found in ALIC_L and PLIC_L. The ALIC carries anterior thalamic radiation and frontopontine fibers, and has been shown to be associated with emotion, decision making, cognition and motivation (35). The PLIC carries fibers of the PTR, CST, and corticopontine tracts, as well as somatosensory fibers from the thalamus to the primary somatosensory cortex, being an important structure for motor, and sensory pathways (35). In this study, we found an interaction between copy-number and age in ALIC, where FA increased with age in deletion carriers, and differed from the typical gradual reduction of FA with age (36), that is seen in NoCNV and duplication carriers (Supplemental Findings and Figure S7). The same effect was also found in BCC and SCC. Although these findings could suggest an atypical development of these regions with age, a younger group (from childhood until adulthood) would be needed to reliably investigate the impact of 15q11.2 BP2-BP2 CNVs on white matter development.

Both samples also showed similar effects (increased FA in deletion carriers) in different portions of the cingulum – significant effects were found on C_CG_L in the Icelandic sample, and on C_HIP_R and C_HIP_L in the UK Biobank sample. The cingulum connects components of the limbic system, where different portions reflect distinct functions (37). The hippocampal portion is linked to learning and episodic memory. Conversely, deletion carriers showed reduced FA and increased RD in the fornix, a major output tract of the hippocampus that is also implicated in memory function. Although not significant, deletion carriers in the Icelandic sample also show a decrease in FA in this structure (Figure S1).

We also found some divergence between the two datasets. While deletion carriers showed increased FA in PTR_R and ILF_L in the Icelandic sample, they showed reduced FA in the UK Biobank sample, with a significant FA reduction in PTR_R when compared to NoCNV carriers. Furthermore, when looking at effect sizes in all tracts (Figure S3), we can see an overall pattern of increased FA in deletion carriers in the Icelandic sample (the only exceptions being fornix and bilateral CST), whereas in the UK Biobank the pattern was more heterogeneous. Effect sizes in both samples are shown for AD, RD and MD in Figures S4-S6. Differences between samples could have resulted from several factors, such as variable gene expression, sample and population characteristics, and different resolution and processing approaches for the imaging data.

NDDs have been generally associated with global decreases in FA (32,38,39), which contrasts with the findings of increased FA in 15q11.2 BP1-BP2 deletion carriers. This raises the question of how changes in FA relate to cognitive function and risk for disorder in these carriers. In our neuroimaging sample, deletion carriers performed worse during reaction time, fluid intelligence, symbol substitution, and trail making B tasks, whereas duplication carriers performed at a similar level as NoCNV controls. The same pattern was observed when extending our analyses to all participants with cognitive data available, where additional effects in pairs matching and digit span tasks were seen in deletion carriers, and the biggest effect size was observed in fluid intelligence (Table S3). This pattern of effects is in line with previous studies, where deletion was reported as being more damaging than duplication for a variety of cognitive tests (6,15).

Our analyses showed that FA in white matter structures affected by carrier status correlate positively with cognitive performance in reaction time, fluid intelligence, symbol substitution, and trail making B tasks, with exception of PLIC_L, which did not correlate with any of the tasks. Mediation analyses revealed that changes in PTR_R partially mediated the effects of the deletion in all cognitive tasks, where the lower FA seen in deletion carriers in this region contributed to 2–6% of the CNV effect across tasks. The PTR is known to connect the caudal parts of the thalamus to both the parietal and occipital lobes (40), and has been previously indicated as the strongest white matter predictor for fluid intelligence (41). On the other hand, the increased FA seen in ALIC_L and C_HIP_L in deletion carriers seemed to have the opposite effect on cognition, removing part of the CNV effect on performance. These findings suggest that increased FA in these regions contribute to better cognitive performance in deletion carriers.

A recent study used data gathered through the Enhancing Imaging Genetic through Meta-Analysis (ENIGMA) consortium (42) and UK Biobank to determine the effects of the 15q11.2 BP1-BP2 CNV on cortical and subcortical brain morphology. The study reported reduced brain surface area and thicker cortex in deletion carriers, where the significant differences in cortical thickness were more evident in the frontal, cingulate and parietal lobes. Furthermore, this study found significant mediation effects of total surface area and cortical thickness on fluid intelligence, with similar proportions as the ones reported in this study (15). Taken together, these findings suggest that 15q11.2 BP1-BP2 CNV effects on white and grey matter provide partially complementary effects on cognitive ability.

Amongst the four genes in this region*, NIPA1* and *CYFIP1* are known to be involved in mechanisms that, when dysregulated, have the potential to alter white matter. *NIPA1* interacts with the bone morphogenic protein (BMP) receptor type II to inhibit BMP signaling, which contributes to axonal growth, guidance and differentiation (43). Enhanced BMP signaling was found to cause abnormal distal axonal overgrowth at the presynaptic neuromuscular junction in a *Drosophila* model (44). *CYFIP1* is considered a likely contributor to 15q11.2 BP1-BP2-associated phenotypes. Dysregulations in this gene result in alterations in dendritic spine morphology and branching (45,46) as well as myelination (47,48). CYFIP1 interacts in two distinct complexes (45): the WAVE regulatory complex, which regulates actin remodeling during neural wiring (49), and the CYFIP1-eIF4E complex, which through interactions with the fragile X mental retardation 1 protein (FMRP) regulates translation of FMRP-target mRNAs (50). FMRP is the gene product of *FMR1,* which when mutated causes fragile X syndrome (FXS), the most common monogenic form of ID (51).

In our previous study, we hypothesized that *CYFIP1* could be a primary contributor to white matter changes in 15q11.2 BP1-BP2 CNV carriers. Previous DTI studies have shown increased FA in FXS patients (52,53), similar to what we observed in 15q11.2 BP1-BP2 deletion carriers, suggesting that these changes could be in part due to disruptions in the CYFIP1-FMRP complex. Recently, we developed a novel *Cyfip1*-haploinsufficient rat line using CRISPR/Cas9 to assess the contribution of *Cyfip1* to white matter microstructure (47). We found evidence that *Cyfip1* can influence myelination, since *Cyfip1* haploinsufficiency led to decreased FA, myelin thinning in the corpus callosum, and aberrant intracellular distribution of myelin basic protein in cultured oligodendrocytes. These findings contrasted with our previous results, from the Icelandic sample, showing widespread increased FA in 15q11.2 BP1-BP2 deletion carriers. However, in the UK Biobank sample, reduced FA was found in PTR_R in deletion carriers, which contributed to worse cognitive performance. Decreased FA could indeed result from myelin deficits (54), that could be caused by dysregulations in *CYFIP1*, which could in turn affect cognition (55,56). Conversely, increased FA in ALIC_L and C_HIP_L led to better cognitive performance in deletion carriers, which could be associated with compensatory mechanisms as a response to primary deficits (e.g. in myelination or synapses). It is, however, difficult to speculate at this point, given that the individual or combined influence of the other three genes in this region on white matter is unknown, and disruptions to myelin and/or axons cannot be distinguished with traditional DTI methods.

The effect sizes reported here were overall smaller than the ones in the Icelandic sample. In this study, we compared carriers to thousands of controls, which provides a better estimate of the general population mean, and therefore a more reliable estimate of effect sizes (57). The deletion has been proposed as a pathogenic CNV of mild effect size (58), which is in line with the smaller effect sizes found in the UK Biobank data. In the Icelandic sample, carriers were compared to 19 NoCNV controls, which may have led to an overestimation of the CNV effect. In addition to the different genetic and environmental background in both samples, participants in the UK Biobank were older and the imaging data were collected and processed differently. Although this could explain some of the variability between the samples, it is encouraging that we found overlapping results from different samples, increasing our confidence of a dosage effect on white matter, particular on the cingulum and internal capsule, which harbor important connections of the limbic system.

It is also important to note that the recruitment in UK Biobank and Icelandic relies on volunteers who put themselves forward to be scanned, which could result in a significant healthy volunteer bias (59). Additionally, in the Icelandic sample, NoCNV controls had similar intelligence quotient scores as carriers. It would then be interesting to compare findings from these two studies with findings from a sample ascertained from clinical services.

### Conclusion

We show converging evidence from two independent samples with different genetic and environmental backgrounds, supporting our hypothesis of a mirrored effect of the 15q11.2 BP1-BP2 CNV on white matter microstructure. We further show that changes in white matter partially mediate the cognitive phenotype in deletion carriers, with decreased FA in the PTR contributing to poorer cognitive performance, while increased FA in ALIC and C_HIP ameliorates, in deletion carriers. This work provides further evidence for gene dosage effects on brain structure and function at the 15q11.2 locus.

## Acknowledgements

This research has been conducted using the UK Biobank Resource under Application Number 17044. This research was supported by the Medical Research Council Programme grant ref. G08005009. This work was also supported by a Wellcome Trust Strategic Award ‘DEFINE’ grant no. 100202/Z/12/Z and core support from the Neuroscience and Mental Health Research Institute, Cardiff University. KMK is supported by a Wellcome Trust Clinical Research Training Fellowship (ref. 201171/Z/16/Z).

