## Supplementary Material for "Analysis of diffusion tensor imaging data from UK Biobank confirms dosage effect of 15q11.2 copy-number variation on white matter and shows association with cognition"

### Supplemental Material

#### Supplemental Methods

##### CNV quality control

The majority of participants in the UK Biobank (~450 000) were genotyped using the Affymetrix UK Biobank Axiom ® array (820 967 probes), with 50 000 participants initially genotyped using the Affymetrix UK BiLEVE Axiom array (807 411 probes). The two arrays were shown to be very similar (over 95% common content). The description of DNA extraction and processing workflow are further described at https://biobank.ctsu.ox.ac.uk/crystal/crystal/docs/genotyping_sample_workflow.pdf. Normalized signal intensity, genotype calls and confidence intervals were generated using ~750 000 biallelic markers that were further processed with PennCNV-Affy software (1). For quality control, individual samples were excluded if they had >30 CNVs, a waviness factor >0.03 or <-0.03, or a call rate <96%. Individual CNVs were also excluded if they were covered by <10 probes or had a density coverage of less than one probe per 20 000 base pairs (2).

##### Details on cognitive data processing

Cognitive data collected from UK Biobank participants at their first assessment at the assessment centres were considered for analyses. When appropriate, data were transformed so that higher scores indicated higher performance.

The Pairs matching task tests episodic memory, where six pairs of cards were shown for three seconds to each participant, then turned over and the participant was asked to identify all correct pairs in as few tries as possible. We used the total number of errors made in round 2 (field 399) of the pairs matching test (field 100030) for analyses, excluding any individual who did not achieve 6 correct matches, in order to exclude participants who did not complete the test. Data were not normally distributed and a log+1 transformation was applied before converting to z scores.

The reaction time task tests simple processing speed, where participants played 12 rounds of the card game 'Snap'. Here, two cards were shown at a time, and the participants had to click a button as fast as possible if the cards were the same. We used the mean time to correctly identify matches (field 20023) for analyses. Outlying scores (1500 ms) were excluded. Data were not normally distributed and a log transformation was applied before converting to z scores.

The Fluid Intelligence Score tests reasoning and problem solving, where thirteen verbal and numerical reasoning questions were made, after which the participants had to complete as many questions as possible within 2 minutes. We used the number of correct answers (field 20016) for analyses. Data were normally distributed, so no transformation was made before converting to z scores.

The Digit Span task tests working memory, where participants were shown a number and asked to recall this once the number had disappeared. The number became one digit longer in each round (total of 12 rounds). We used maximum number of digits correctly recalled (field 4282) for analyses. Data were normally distributed, so no transformation was made before converting to z scores.

The Symbol Digit Substitution task tests complex processing speed, where participants were asked to play a code-breaking game, which consisted in matching numbers to a set of symbols within 2 minutes. We used the number of symbol digit matches made correctly (field 20159) for analyses. Outlying scores (0-1 and >36) were excluded. Data were normally distributed, after outlier exclusion, so no transformation was made before converting to z scores.

The Trail Making A and B tasks test visual attention and consists of both a numeric (A) and an alphanumeric (B) part. Participants were asked to connect circles with numbers (trail A), and with alternating numbers and letters (trail B), in the correct ascending order. For both tests, we used the time to complete the task (trail A – field 20156, trail B – field 20157) for analyses. Data were not normally distributed, and a log transformation was applied before converting to z scores.

### Supplemental Finding

#### Age and sex interaction

We found significant interactions between copy-number and age on FA in BCC, SCC, ALIC_R, and ALIC_L (Table S4). Age trajectory plots, shown in Figure S7, suggest that this interaction is driven by deletion carriers, who do not follow the same age trajectory as NoCNV- and duplication carriers in which FA progressively decreases with age. Instead, deletion carriers seem to maintain or increase FA values with age. This pattern does not extend to cognitive performance, where no interactions between copy-number and age were found (Table S5 and Figure S8). We found an interaction between copy-number and sex only on FA in C_CG_R (Table S6). No significant interaction between copy-number and sex was observed regarding cognitive performance (Table S7).

**Supplemental Table 1** – Critical region coordinates, calling criteria, and number of genes hit for the 15q11.2 BP1-BP2 copy number variant in the UK Biobank Sample.

| CNV | Locus | **Critical/Unique Sequence Region (hg19)** | **N Biallelelic Axiom probes** | **PSD Genes (from 685 PSD genes)** | **Calling criteria** | **N genes** |
|---|---|---|---|---|---|---|
| 15q11.2 BP1-BP2 | 15q11.2 | chr15:22,805,313-23,094,530 | 145 | *CYFIP1* | Size >50% of critical region | 5 |

**Supplemental Table 2** – Effects of the15q11.2 BP1-BP2 copy number variant on TBSS-derived measures. Results from TukeyHSD pairwise comparisons are shown where both uncorrected p-values (unc.) and FDR corrected p-values (BH corr.) are represented, as well as Cohen's d effect sizes for each comparison. Only white matter tracts showing any significant group differences after FDR correction are shown.

| Dependent variable | Contrast | ROI | t value | p-value (unc.) | p-value (BH corr.) | Effect size Cohen's d (SE) |
|---|---|---|---|---|---|---|
| FA | Del vs NoCNV | fornix | -2.77 | 0.006 | 0.06 | -0.28 (0.20) |
| | | ALIC_L | 3.12 | 0.002 | **0.03** | 0.31 (0.19) |
| | | PLIC_R | 2.08 | 0.04 | 0.16 | 0.21 (0.19) |
| | | PLIC_L | 3.11 | 0.002 | **0.03** | 0.31 (0.19) |
| | | PTR_R | -3.39 | 0.0007 | **0.02** | -0.33 (0.19) |
| | | C_HIP_R | 2.19 | 0.03 | 0.13 | 0.22 (0.20) |
| | | C_HIP_L | 3.96 | 0.00008 | **0.007** | 0.39 (0.20) |
| | Dup vs NoCNV | fornix | 2.25 | 0.02 | 0.12 | 0.21 (0.18) |
| | | ALIC_L | -1.40 | 0.2 | 0.40 | -0.13 (0.18) |
| | | PLIC_R | -2.22 | 0.03 | 0.13 | -0.20 (0.18) |
| | | PLIC_L | -1.71 | 0.09 | 0.27 | -0.16 (0.18) |
| | | PTR_R | 0.67 | 0.5 | 0.8 | 0.06 (0.18) |
| | | C_HIP_R | -3.19 | 0.001 | **0.02** | -0.29 (0.18) |
| | | C_HIP_L | -2.26 | 0.02 | 0.12 | -0.21 (0.18) |
| | Del vs Dup | fornix | -3.57 | 0.0004 | **0.01** | -0.49 (0.27) |
| | | ALIC_L | 3.24 | 0.001 | **0.03** | 0.43 (0.27) |
| | | PLIC_R | 3.04 | 0.002 | **0.04** | 0.42 (0.27) |

| | | | | | | |
|---|---|---|---|---|---|---|
| | | PLIC_L | 3.45 | 0.0006 | **0.02** | 0.49 (0.27) |
| | | PTR_R | -2.93 | 0.003 | **0.04** | -0.39 (0.27) |
| | | C_HIP_R | 3.78 | 0.0002 | **0.01** | 0.52 (0.27) |
| | | C_HIP_L | 4.43 | 9.32e-06 | **0.003** | 0.61 (0.28) |
| MD | Del vs NoCNV | BCC | -3.22 | 0.001 | **0.03** | -0.32 (0.20) |
| | | fornix | 2.90 | 0.004 | **0.05** | 0.29 (0.20) |
| | | UF_L | -3.68 | 0.0002 | **0.01** | -0.37 (0.20) |
| | Dup vs NoCNV | BCC | 1.27 | 0.20 | 0.5 | 0.12 (0.18) |
| | | fornix | -3.00 | 0.003 | 0.04 | -0.28 (0.18) |
| | | UF_L | -0.69 | 0.58 | 0.76 | -0.06 (0.18) |
| | Dup vs NoCNV | BCC | -3.23 | 0.001 | **0.03** | -0.48 (0.27) |
| | | fornix | 4.18 | 2.99e-05 | **0.004** | 0.58 (0.27) |
| | | UF_L | -2.21 | 0.03 | 0.13 | -0.33 (0.27) |
| AD | Del vs NoCNV | BCC | -3.76 | 0.0002 | **0.01** | -0.37 (0.19) |
| | | SCC | -3.56 | 0.0004 | **0.01** | -0.35 (0.19) |
| | | fornix | 2.30 | 0.02 | 0.1 | 0.23 (0.20) |
| | | PLIC_L | 3.07 | 0.002 | **0.03** | 0.30 (0.19) |
| | Dup vs NoCNV | BCC | -0.14 | 0.9 | 0.9 | -0.01 (0.18) |
| | | SCC | -0.50 | 0.62 | 0.8 | -0.05 (0.18) |
| | | fornix | -2.73 | 0.006 | 0.06 | -0.26 (0.19) |
| | | PLIC_L | -0.63 | 0.53 | 0.78 | -0.06 (0.18) |
| | Del vs Dup | BCC | -2.65 | 0.008 | 0.07 | -0.39 (0.27) |
| | | SCC | -2.26 | 0.02 | 0.1 | -0.34 (0.27) |
| | | fornix | 3.56 | 0.0004 | **0.01** | 0.52 (0.28) |
| | | PLIC_L | 2.67 | 0.008 | 0.07 | 0.38 (0.27) |
| RD | | fornix | 3.09 | 0.002 | **0.03** | 0.31 (0.20) |

| | | | | | | |
|---|---|---|---|---|---|---|
| | Del vs NoCNV | PTR_R | 3.40 | 0.0007 | **0.02** | 0.34 (0.19) |
| | | C_HIP_R | -2.77 | 0.006 | 0.06 | -0.06 (0.19) |
| | Dup vs NoCNV | fornix | -2.74 | 0.006 | 0.06 | -0.26 (0.18) |
| | | PTR_R | -1.73 | 0.24 | 0.51 | -0.11 (0.18) |
| | | C_HIP_R | 2.37 | 0.02 | 0.11 | 0.19 (0.18) |
| | Del vs Dup | fornix | 4.14 | 3.53e-05 | **0.004** | 0.57 (0.27) |
| | | PTR_R | 3.29 | 0.001 | **0.02** | 0.42 (0.27) |
| | | C_HIP_R | -3.65 | 0.0003 | **0.01** | -0.26 (0.27) |

**Supplemental Table 3 –** Effects of the 15q11.2 BP1-BP2 copy number variant on seven cognitive tasks from the UK Biobank study. All individuals with cognitive data available were used in this analysis. Group differences were assessed using an ANOVA followed by *post hoc* pairwise comparisons, and corrected using FDR correction. Both uncorrected and corrected (BH corr.) p-values are shown.

**Abbreviations used:** Del, deletion; Dup, duplication; NoCNV, no large copy number variants.

| | 15q11.2 BP1-BP2 CNV | | | Deletion vs NoCNV | | | Duplication vs NoCNV | | | Dosage | |
|---|---|---|---|---|---|---|---|---|---|---|---|
| | Del, No. | NoCNV, No. | Dup, No. | Cohen's d (SE) | p-value | p-value (BH corr.) | Cohen's d (SE) | p-value | p-value (BH corr.) | F statistic | p-value |
| Pairs matching | 1472 | 362,646 | 1789 | -0.11 (0.05) | **1.32e-05** | **4.62e-05** | -0.06 (0.05) | 0.009 | **0.02** | $F_{(2,365899)}$ = 12.13 | 5.38e-06 |
| Reaction time | 1507 | 368,189 | 1820 | -0.23 (0.05) | **< 2e-16** | **4.2e-15** | -0.03 (0.05) | 0.2 | 0.27 | $F_{(2,371508)}$ = 36.59 | <2e-16 |
| Fluid Intelligence | 456 | 119,040 | 582 | -0.34 | **5.21e-13** | **5.47e-12** | 0.04 (0.08) | 0.3 | 0.35 | $F_{(2,120070)}$ = 25.72 | 6.79e-12 |
| Digit Span | 97 | 26,227 | 107 | -0.27 (0.19) | 0.007 | **0.02** | -0.16 (0.19) | 0.09 | 0.16 | $F_{(2,26423)}$ = 4.42 | 0.01 |
| Symbol substitution | 340 | 93,128 | 391 | -0.17 (0.11) | **0.002** | **0.005** | -0.01 (0.1) | 0.8 | 0.8 | $F_{(2,93851)}$ = 6.02 | 0.002 |
| Trail making A | 302 | 82,349 | 342 | -0.09 (0.11) | 0.120 | 0.19 | -0.07 (0.11) | 0.2 | 0.27 | $F_{(2,82985)}$ = 2.37 | 0.09 |
| Trail making B | 302 | 82,348 | 342 | -0.28 (0.11) | **9.43e-07** | **3.96e-06** | -0.03 (0.11) | 0.6 | 0.7 | $F_{(2,82984)}$ = 13.26 | 1.75e-06 |

**Supplemental Table 4 –** Results for fractional anisotropy, including an interaction term between copy number and age. The p-values shown are not corrected.

| | Copy number | | Age | | Copy number * Age | |
|---|---|---|---|---|---|---|
| WT tract | F statistics | p-value | F statistics | p-value | F statistics | p-value |
| BCC | $F_{(2,29624)}=2.11$ | 0.12 | $F_{(1,29624)}=2326.23$ | < 2e-16 | $F_{(2,29624)}=3.99$ | **0.019** |
| SCC | $F_{(2,29690)}=0.84$ | 0.43 | $F_{(1,29690)}=97.98$ | <2e-16 | $F_{(1,29690)}=3.30$ | **0.037** |
| ALIC_R | $F_{(2,29464)}=3.67$ | 0.026 | $F_{(1,29464)}=1050.89$ | <2e-16 | $F_{(2,29464)}=3.67$ | **0.025** |
| ALIC_L | $F_{(2,29327)}=6.57$ | 0.001 | $F_{(1,29327)}=973.32$ | <2e-16 | $F_{(2,29327)}=3.23$ | **0.039** |

**Supplemental Table 5 –** Results for cognitive performance, including an interaction term between copy number and age. All individuals with cognitive data available were used in this analysis. The p-values shown are not corrected.

| | Copy number | | Age | | Copy number * Age | |
|---|---|---|---|---|---|---|
| Test | F statistics | p-value | F statistics | p-value | F statistics | p-value |
| Pairs matching | $F_{(2,365897)}=12.13$ | 5.38e-06 | $F_{(1,365897)}=10904.58$ | < 2e-16 | $F_{(2,365897)}=2.33$ | 0.1 |
| Reaction time | $F_{(2,371506)}=36.59$ | <2e-16 | $F_{(1,371506)}=44113.94$ | < 2e-16 | $F_{(2,371506)}=0.96$ | 0.38 |
| Fluid Intelligence | $F_{(2,120068)}=25.72$ | 6.79e-12 | $F_{(1,120068)}=1004.19$ | < 2e-16 | $F_{(2,120068)}=2.41$ | 0.09 |
| Digit Span | $F_{(2,26421)}=4.42$ | 0.01 | $F_{(1,26421)}=600.29$ | < 2e-16 | $F_{(2,26421)}=0.28$ | 0.76 |
| Symbol substitution | $F_{(2,93849)}=6.02$ | 0.002 | $F_{(1,93849)}=23501.77$ | < 2e-16 | $F_{(2,93849)}=1.09$ | 0.33 |
| Trail making A | $F_{(2,82983)}=2.37$ | 0.09 | $F_{(2,82983)}=9325.07$ | < 2e-16 | $F_{(2,82983)}=2.62$ | 0.07 |
| Trail making B | $F_{(2,82982)}=13.26$ | 1.75e-06 | $F_{(2,82982)}=15183.41$ | < 2e-16 | $F_{(2,82982)}=0.35$ | 0.71 |

**Supplemental Table 6 –** Results for fractional anisotropy, including an interaction term between copy number and sex. The p-values shown are not corrected.

| | Copy number | | Sex | | Copy number * Sex | |
|---|---|---|---|---|---|---|
| WT tract | F statistics | p-value | F statistics | p-value | F statistics | p-value |
| C_CG_R | $F_{(2,29646)}$ =4.71 | 0.009 | $F_{(1, 29646)}$ =401.72 | < 2e-16 | $F_{(2, 29646)}$ =4.08 | **0.017** |

**Supplemental Table 7 –** Results for cognitive performance, including an interaction term between copy number and sex. All individuals with cognitive data available were used in this analysis. The p-values shown are not corrected.

| | Copy number | | Sex | | Copy number * Sex | |
|---|---|---|---|---|---|---|
| Test | F statistics | p-value | F statistics | p-value | F statistics | p-value |
| Pairs matching | $F_{(2,365897)}$ =12.13 | 5.38e-06 | $F_{(1,365897)}$ = 98.09 | < 2e-16 | $F_{(2,365897)}$ =0.46 | 0.63 |
| Reaction time | $F_{(2,371506)}$ = 36.59 | <2e-16 | $F_{(1,371506)}$ = 3281.89 | < 2e-16 | $F_{(2,371506)}$ = 0.57 | 0.57 |
| Fluid Intelligence | $F_{(2,120068)}$ = 25.72 | 6.79e-12 | $F_{(1,120068)}$ = 441.62 | < 2e-16 | $F_{(2,120068)}$ = 0.24 | 0.79 |
| Digit Span | $F_{(2,26421)}$ = 4.42 | 0.01 | $F_{(1,26421)}$ = 198.97 | < 2e-16 | $F_{(2,26421)}$ = 0.1 | 0.91 |
| Symbol substitution | $F_{(2,93849)}$ = 6.02 | 0.002 | $F_{(1,93849)}$ = 97.59 | < 2e-16 | $F_{(2,93849)}$ = 0.34 | 0.71 |
| Trail making A | $F_{(2,82983)}$ = 2.37 | 0.09 | $F_{(2,82983)}$ = 342.23 | < 2e-16 | $F_{(2,82983)}$ = 0.39 | 0.68 |
| Trail making B | $F_{(2,82982)}$ = 13.26 | 1.75e-06 | $F_{(2,82982)}$ = 76.56 | < 2e-16 | $F_{(2,82982)}$ = 0.26 | 0.77 |

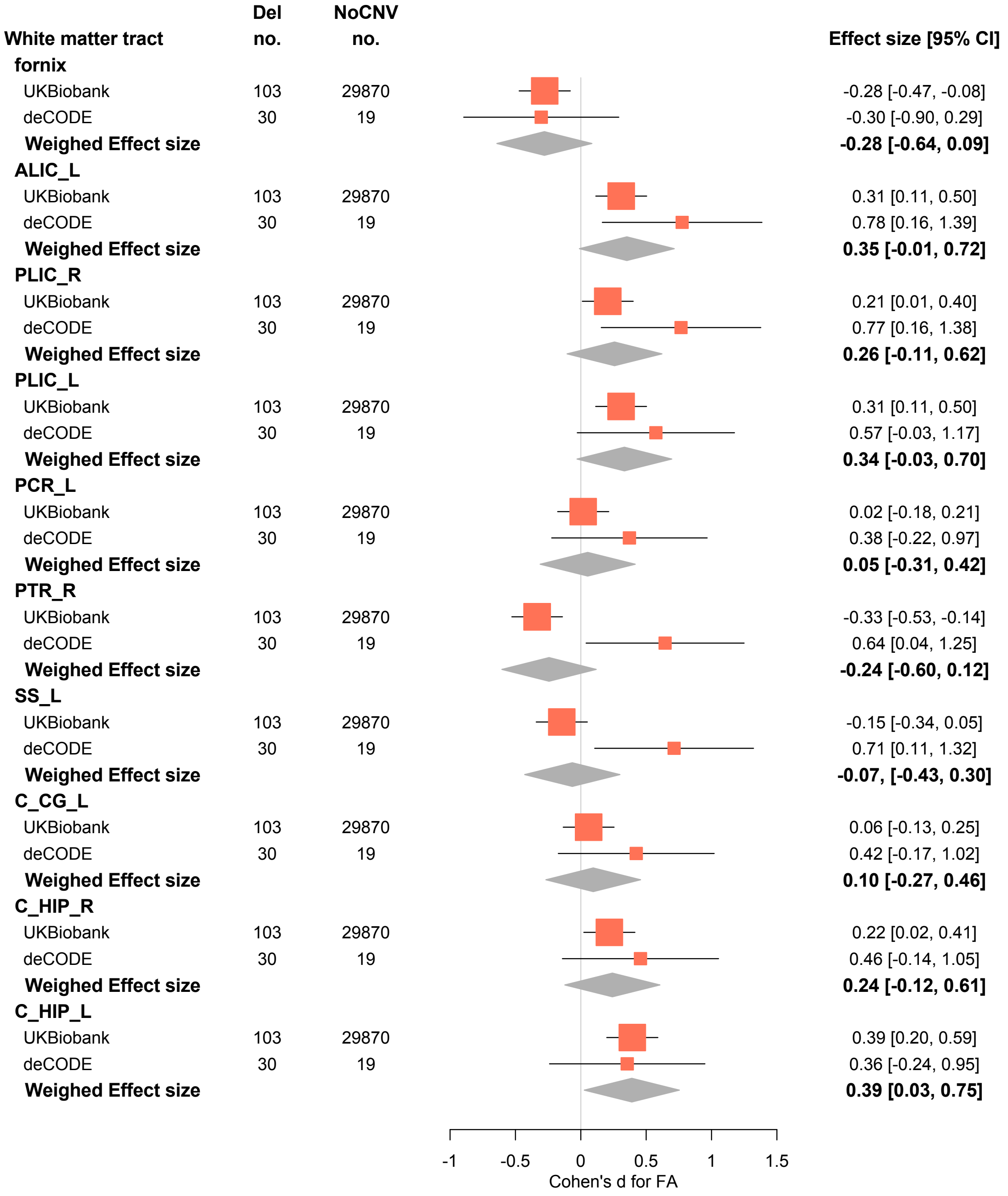


**Supplementary Figure 1 -** Forest plot showing the Cohen's d effect sizes in the UK Biobank and the Icelandic sample, relative to the comparison between deletion (Del) and NoCNV carriers on fractional anisotropy. Here, only white matter tracts showing group effects in the UKBiobank sample and/or in the Icelandic sample are shown. The summary diamond shows the weighed effect size of both studies, calculated using the inverse of the variance method.

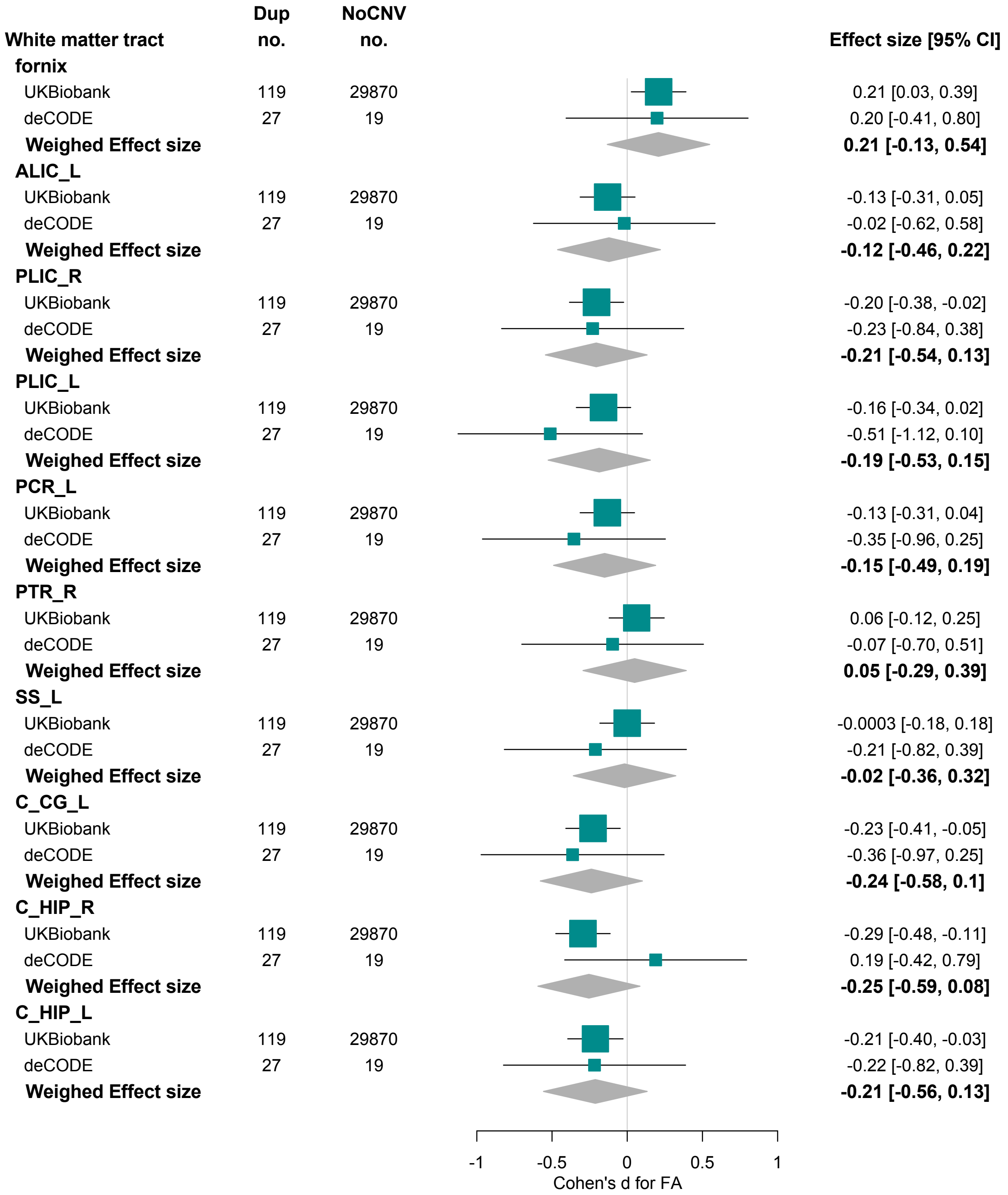


**Supplementary Figure 2** -- Forest plot showing the Cohen's d effect sizes in the UK Biobank and the Icelandic sample, relative to the comparison between duplication (Dup) and NoCNV carriers on fractional anisotropy. Here, only white matter tracts showing group effects in the UKBiobank sample and/or in the Icelandic sample are shown. The summary diamond shows the weighed effect size of both studies, calculated using the inverse of the variance method.

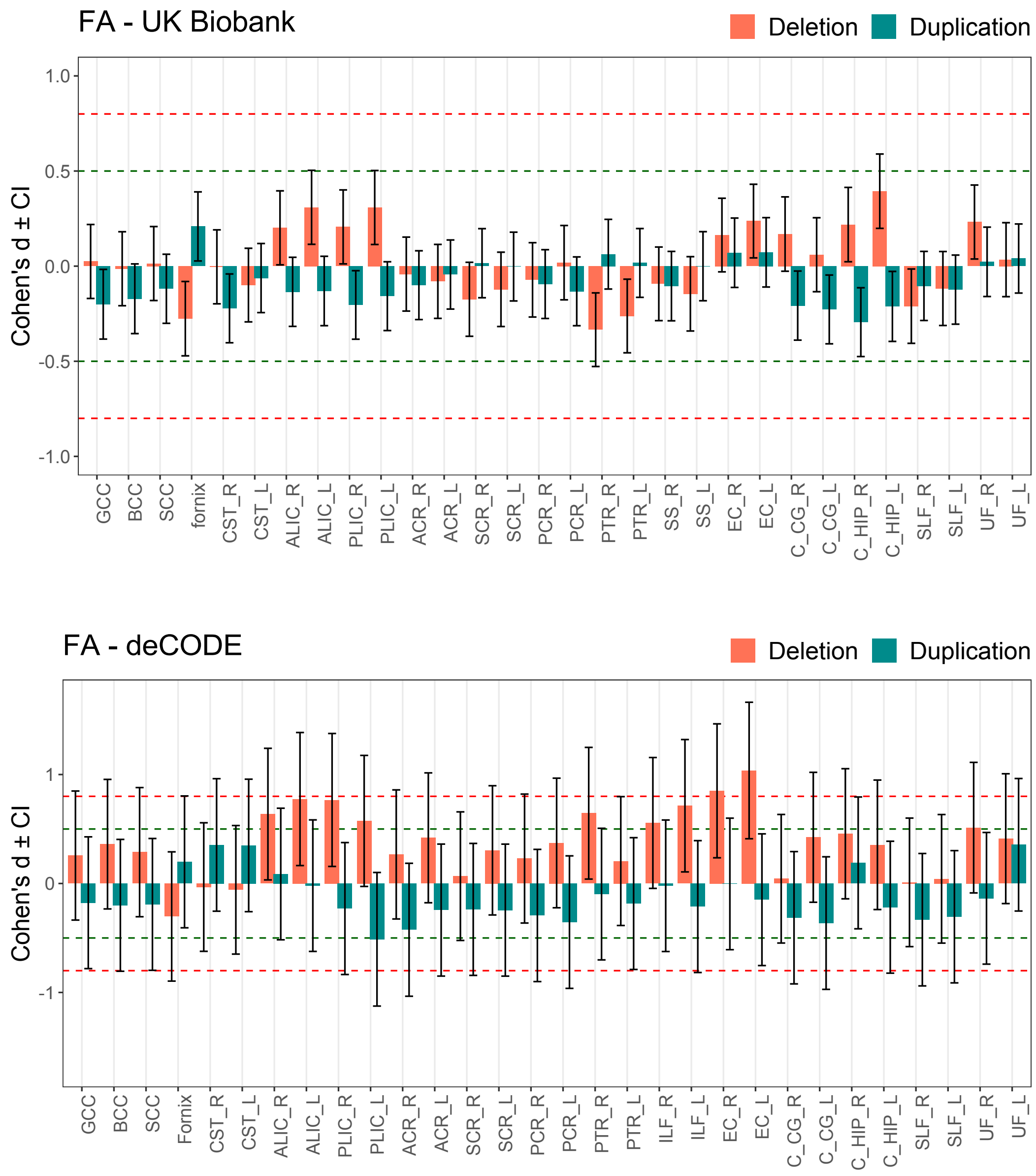


**Supplemental Figure 3** – Diverging bars showing Cohen's d effect sizes for fractional anisotropy in the UK Biobank sample (top) and in the Icelandic sample (bottom), for the 30 white matter tracts considered in the study. In the UK Biobank sample, 103 deletion and 119 duplication carriers were compared to 29870 NoCNV carriers. In the Icelandic sample, 30 deletion and 27 duplication carriers were compared to 19 NoCNV carriers. The thresholds where effects sizes are considered large (0.8) or medium (0.5), according to Cohen's criteria, are represented by a vertical red or green dashed line, respectively.

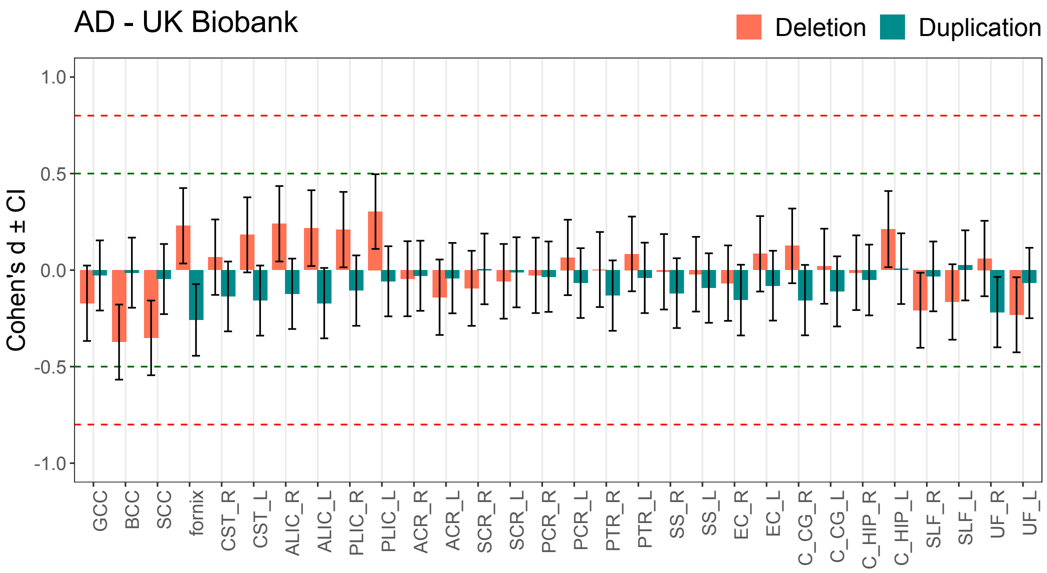


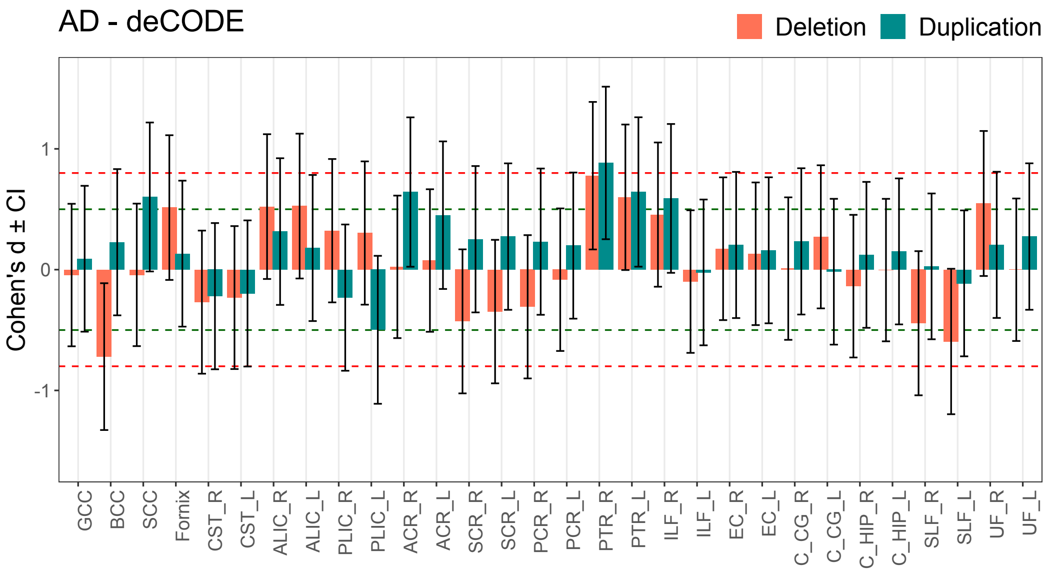


**Supplemental Figure 4** – Diverging bars showing Cohen's d effect sizes for axial diffusivity in the UK Biobank sample (top) and in the Icelandic sample (bottom), for the 30 white matter tracts considered in the study. In the UK Biobank sample, 103 deletion and 119 duplication carriers were compared to 29870 NoCNV carriers. In the Icelandic sample, 30 deletion and 27 duplication carriers were compared to 19 NoCNV carriers. The thresholds where effects sizes are considered large (0.8) or medium (0.5), according to Cohen's criteria, are represented by a vertical red or green dashed line, respectively.

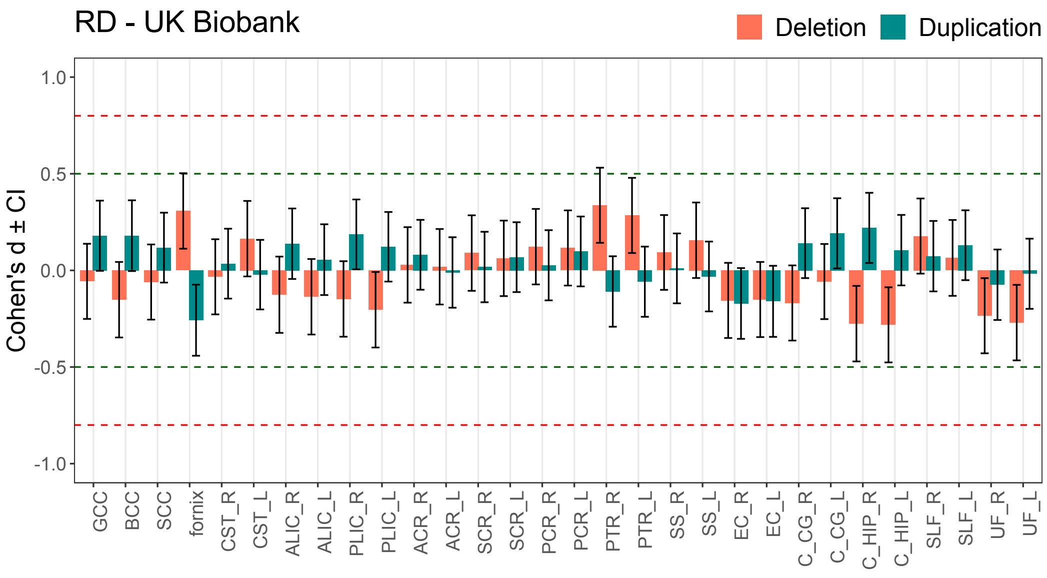


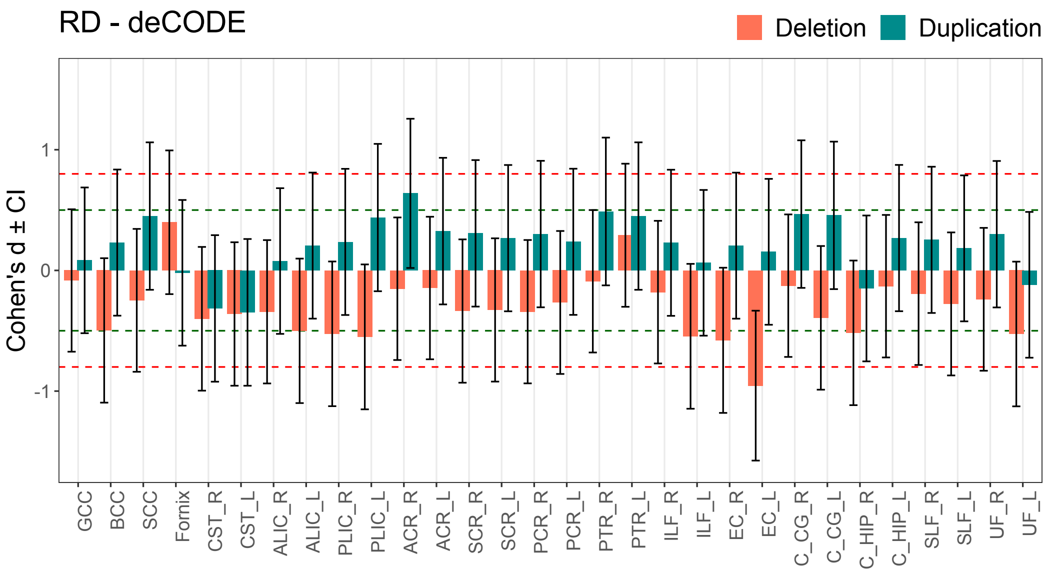


**Supplemental Figure 5** - Diverging bars showing Cohen's d effect sizes for radial diffusivity in the UK Biobank sample (top) and in the Icelandic sample (bottom), for the 30 white matter tracts considered in the study. In the UK Biobank sample, 103 deletion and 119 duplication carriers were compared to 29870 NoCNV carriers. In the Icelandic sample, 30 deletion and 27 duplication carriers were compared to 19 NoCNV carriers. The thresholds where effects sizes are considered large (0.8) or medium (0.5), according to Cohen's criteria, are represented by a vertical red or green dashed line, respectively.

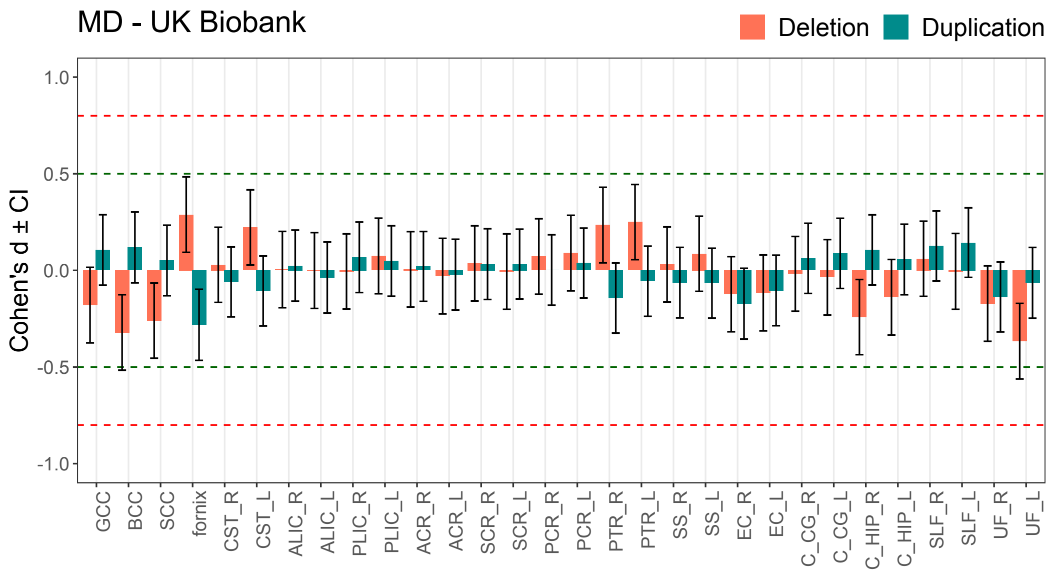


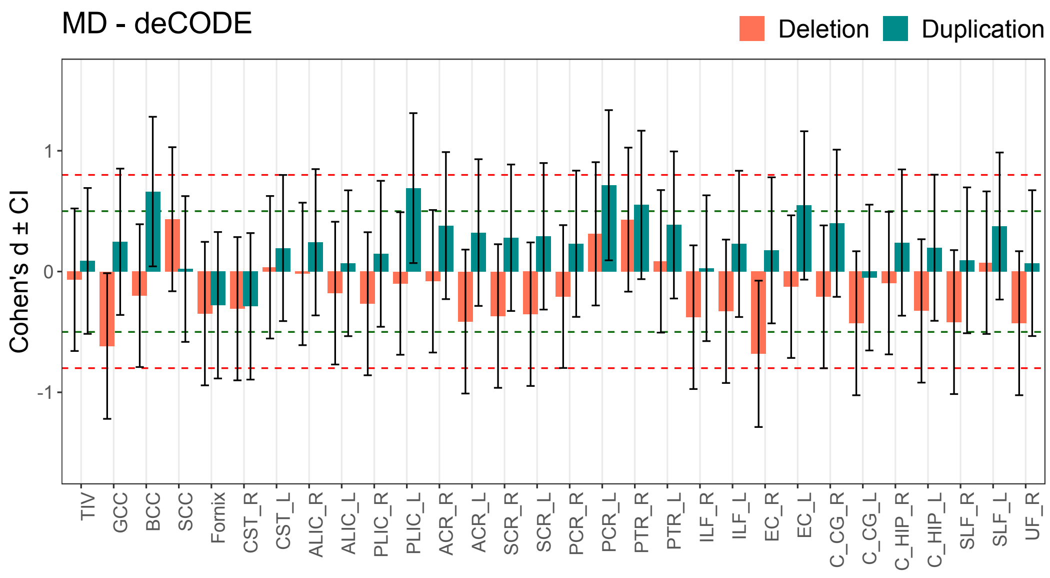


**Supplemental Figure 6** - Diverging bars showing Cohen's d effect sizes for mean diffusivity in the UK Biobank sample (top) and in the Icelandic sample (bottom), for the 30 white matter tracts considered in the study. In the UK Biobank sample, 103 deletion and 119 duplication carriers were compared to 29870 NoCNV carriers. In the Icelandic sample, 30 deletion and 27 duplication carriers were compared to 19 NoCNV carriers. The thresholds where effects sizes are considered large (0.8) or medium (0.5), according to Cohen's criteria, are represented by a vertical red or green dashed line, respectively.

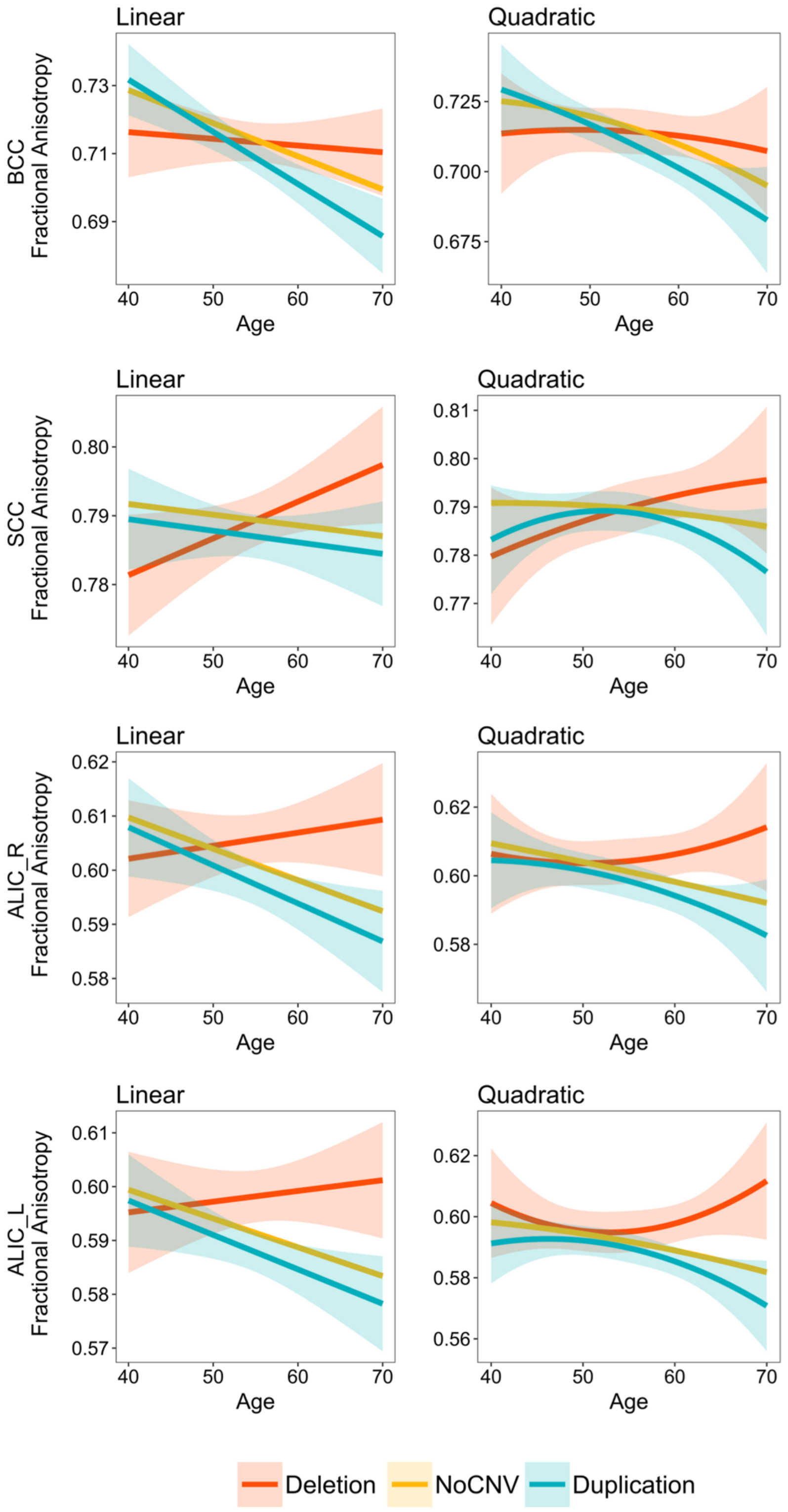


**Supplemental Figure 7 –** Age trajectories showing the linear (left) and quadratic (right) relationship between fractional anisotropy and age. The three regression lines shown represent the three carrier groups, as indicated in the legend.

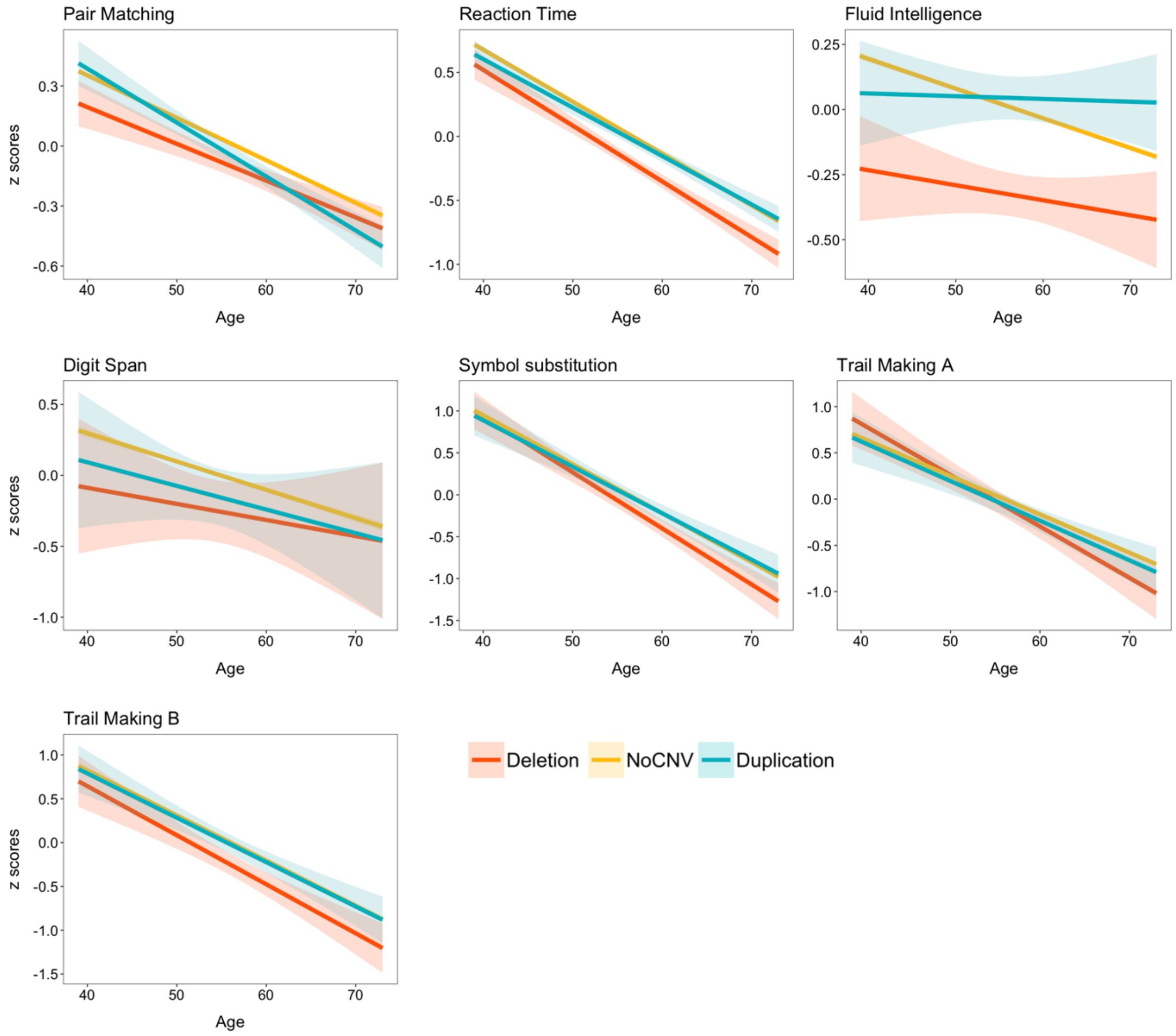


**Supplemental Figure 8** – Age trajectories showing the linear relationship between cognitive performance for each test and age. The three regression lines shown represent the three carrier groups, as indicated in the legend.
